# Defining the transmission parameters of lumpy skin disease virus by the three model insect vector species *Aedes aegypti*, *Stomoxys calcitrans* and *Culicoides nubeculosus* reveals important differences in their likely role in transmission

**DOI:** 10.1101/2025.10.11.681786

**Authors:** Henry Munyanduki, Beatriz Sanz Bernardo, Najith Wijesiriwardana, Ismar R. Haga, Petra C. Fay, Charlotte G. Cook, Isabel Bacon, Alice Catanzaro, Christopher J. Sanders, John Atkinson, Carrie Batten, Luke Alphey, Karin E Darpel, Simon Gubbins, Philippa M. Beard

**Affiliations:** The Pirbright Institute, Ash Rd, Pirbright, GU24 0NF, UK; MSD Animal Health, Walton Manor, Walton, Milton Keynes MK7 7AJ, UK; The Department of Biology, University of York, Wentworth Way, York, YO10 5DD, UK; York Biomedical Research Institute, University of York, Heslington, YO10 5DD, UK; Keele University, Keele, Staffordshire, ST5 5BG, UK; Institute of Virology and Immunology, Mittelhäusern, Switzerland; Department of Infectious Diseases and Pathobiology, Vetsuisse Faculty, University of Bern, Bern, Switzerland; Australian Centre for Disease Preparedness CSIRO, 5 Portalington Road, Geelong, Victoria, 3220, Australia; Hartpury University and Hartpury College, Hartpury, GL19 3BE, UK

**Keywords:** lumpy skin disease, poxvirus, insect vector, bovine

## Abstract

Lumpy skin disease virus (LSDV) is a poxvirus that can cause severe, systemic disease in cattle. By far the most important route of transmission of LSDV is mechanical transmission via haematophagous arthropod vectors. However we lack detailed information on this process including the likelihood of transmission by different vector species. This study used an experimental bovine model of LSDV transmission to quantify the transmission of LSDV from an infected donor to a naïve recipient calf. Three species of Diptera representing different vector groups were included (*Stomoxys calcitrans*, *Aedes aegypti* and *Culicoides nubeculosus,* respectively a large biting fly, a mosquito and a midge), and the clinical, virological and immunological outcomes in the recipient calves studied. The ability of *Ae. aegypti* to mechanically transmit LSDV following feeding on an artificial membrane feeding system was also examined. Both *Ae. aegypti* and *S. calcitrans* were able to transmit LSDV, resulting in disease in recipient calves. Bites from virus-positive *C. nubeculosus* did not result in disease in recipient calves, though the presence of neutralising antibodies in these recipients indicated exposure to virus or virus components. *Ae. aegypti* successfully transmitted LSDV following feeding on LSDV-spiked blood through an artificial membrane feeding system, validating this laboratory model as a future replacement for donor cattle. Mathematical models of the data were generated and predicted *S. calcitrans* to be the most efficient vector of LSDV of the insects tested with a reproduction number (R_0_) of 5.8.

**Impact statement:** Lumpy skin disease virus (LSDV) is a neglected, rapidly emerging pathogen of cattle that has spread into Europe and throughout the Middle East and Asia over the past ten years. Lack of understanding of the mechanism of transmission of LSDV has hampered efforts to control its rapid spread. This study compares the ability of three model species of Diptera (*Stomoxys calcitrans*, *Aedes aegypti* and *Culicoides nubeculosus*) to mechanically transmit the virus to cattle, generating high quality quantitative data to facilitate mathematical modelling of virus transmission. *S. calcitrans* was identified as a potential driver of LSDV transmission with a *R*_0_ of 5.8. This work provides new insights into the vector transmission of LSDV that can be used to design more targeted and effective disease control programmes.

## Introduction

Lumpy skin disease (LSD) is a viral disease affecting cattle, caused by infection with the poxvirus lumpy skin disease virus (LSDV). Clinically infected animals are characterised by the presence of raised multifocal cutaneous skin lesions, weight loss, reproductive failure, and death (Sanz-Bernardo et al., 2020). The disease results in poor animal welfare, economic losses, and trade barriers. In the past decade LSD has spread from Africa and the Middle East into Europe and Asia causing substantial loss to affected regions (Bianchini et al., 2023, Wilhelm and Ward, 2023, Hunter and Wallace, 2001, Tuppurainen et al., 2017, Beard, 2016, Carn and Kitching, 1995a, Lee et al., 2024). More recently LSD has been reported for the first time in both Italy and France (DEFRA, 2025). An incomplete understanding of the mechanism of transmission of LSDV has hampered the design and implementation of control and eradication programmes.

Experimental bovine studies by Carn and Kitching (1995) demonstrated an absence of direct transmission of LSDV from infected to naïve cattle housed together (Carn and Kitching, 1995b). In the same study the minimum dose of virus required to cause generalised disease by intradermal inoculation was shown to be 10^2^ TCID_50_/ml (Carn and Kitching, 1995a). This study, coupled with field observations that LSD outbreaks occurred during wet seasons when insect numbers were high, led researchers to hypothesise the role of biting insects in LSDV transmission (Gari et al., 2010, Burdin and Prydie, 1959, Kahana-Sutin et al., 2017, Magori-Cohen et al., 2012).

This hypothesis was supported by the knowledge that other species of poxvirus are transmitted via haematophagous arthropods. Experimental studies of the leporipoxviruses myxoma virus and shope fibroma virus have shown that these viruses are spread between rabbits by mosquitoes. The evidence was consistent with the mosquito mouthparts becoming contaminated with virus after feeding on affected rabbits ((Filshie, 1964) and reviewed by (Dalmat and Cunningham, 1959)). No replication of the virus was observed in these insects suggesting that they were acting as mechanical rather than biological vectors (Day et al., 1956). Similar findings have been reported when describing transmission of fowlpox virus (Kligler et al., 1929).

Insect mediated transmission of LSDV was first demonstrated under experimental conditions by Chihota and colleagues. A group of *Aedes aegypti* mosquitoes that had fed on clinically-affected donor cattle transmitted LSDV after sequential feeding on 7 susceptible naïve cattle. Five of the seven animals developed LSD with clinical signs similar to natural LSD (Chihota et al., 2001). This work was used to model the risk of vector transmission of LSDV (Gubbins, 2019).

More recent work has shown that the stable fly *S. calcitrans* and *Tabanidae* horseflies *Haematopota* spp can mechanically transmit LSDV from clinical to naïve susceptible animals (Sohier et al., 2019, Haegeman et al., 2023). A similar experimental study found evidence of LSDV transmission from clinical donor to naive recipient animal by three *Stomoxys* species (*S. calictrans, S. sitiens* and *S. indica*) (Issimov et al., 2020). A limitation of these studies was that the quantities of insects that fed on the animals that became clinical could not be determined.

We have published studies on the initial stages of LSDV transmission to estimate LSDV acquisition and retention by four insect species (*Ae. aegypti*, *Culex quinquefasciatus*, *S. calcitrans*, and *C. nubeculosus*). Insects that fed on clinically infected animals were individually analysed, revealing that all four insect species readily acquired LSDV from infected cattle. No propagation of LSDV was detected in the insects, consistent with the mode of mechanical mechanism of transmission. The highest titres of LSDV were found in the skin lesions compared to normal skin of an infected animal. Consistent with this finding LSDV was readily acquired by the vectors feeding on skin lesions, with acquisition from normal skin uncommon (Sanz-Bernardo et al., 2021). Virus was found to be preferentially retained in the proboscis and head for up to 8 days (Sanz-Bernardo et al., 2022), consistent with previous work on the leporipoxvirus myxoma virus (Filshie, 1964).

These studies examined the acquisition and retention of LSDV in a range of potential insect vectors. However, the limited data available for LSDV transmission by different vector species, particularly uncertainty regarding the efficiency of LSDV transmission by different vector species, was highlighted as a key knowledge gap (Gubbins, 2019, Sanz-Bernardo et al., 2021). To address this we carried out a series of *in vivo* experiments with three model species representing different vector groups; the mosquito *Ae. aegypti*, the large biting fly *S. calcitrans*, and the biting midge *C. nubeculosus*, and assessed their ability to transmit LSDV between cattle. The data from the resulting transmission parameters was used to calculate the *R*_0_ of the different insect vectors. This has revealed markedly different transmission parameters for different insect species, suggesting the vector groups play different roles in the transmission of LSDV in the field.

## Results

Five experiments (studies A-E) were carried out to quantify the transmission of LSDV between cattle by insect vectors. The studies were based around transmission of LSDV from a clinically-affected “donor” calf or an artificial feeding membrane system to a naïve “recipient” calf, as summarised in **Figure 1** and **Table 1**. The transmission potential of three model insect species (*Ae. aegypti* mosquitoes, *S. calcitrans* flies, and *C. nubeculosus* midges) was studied. Clinical, virological, and immunological data were collected from each recipient calf to characterise in detail the disease resulting from vector transmission of LSDV. A key design feature of the experiments was the analysis of each individual insect enabling us to generate quantitative data that was then used to model LSDV transmission and calculate the risk of transmission associated with each insect species.

**Figure 1:**
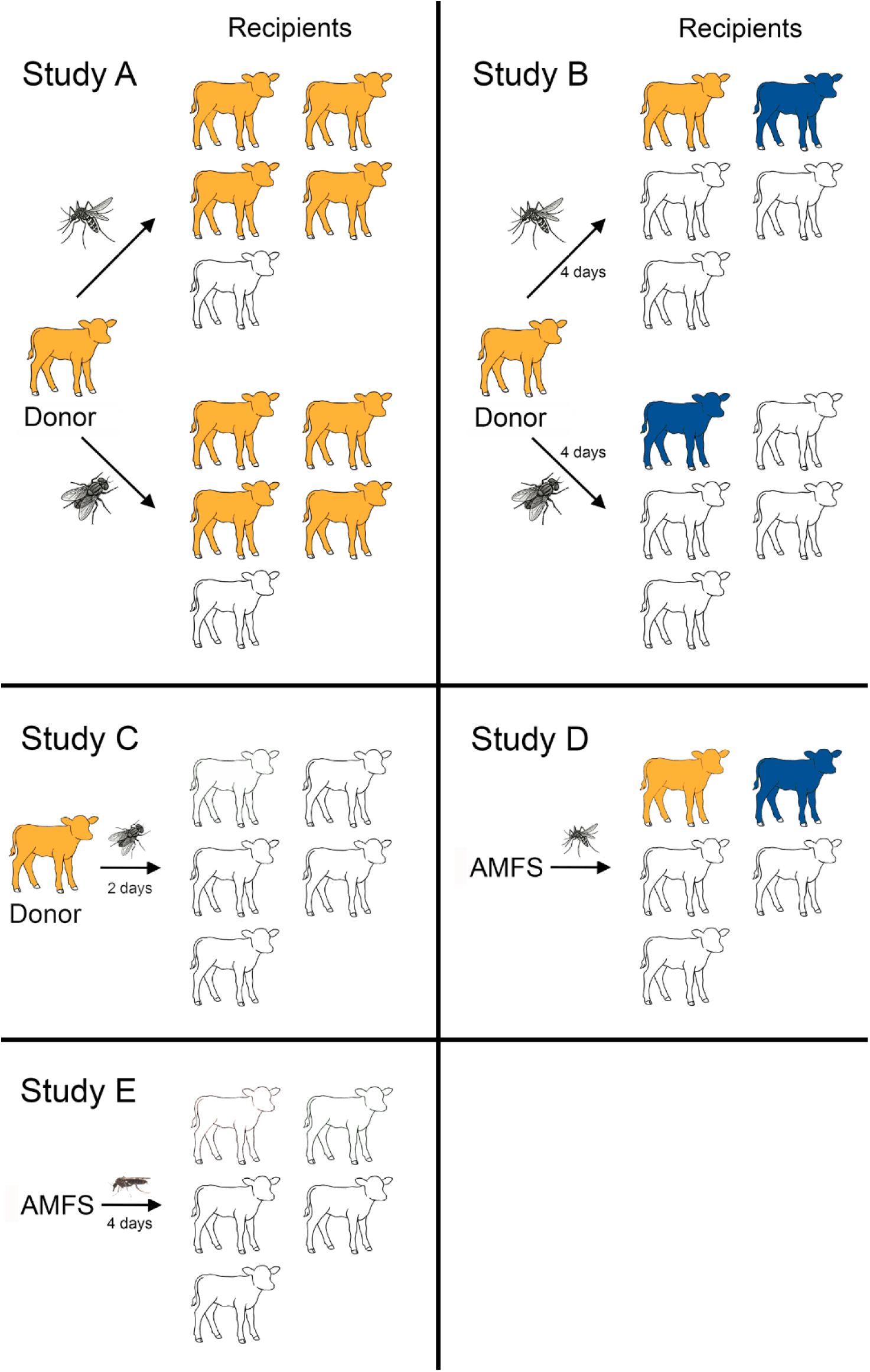
Infographic summarising the design of the five studies (A-E) and key outcomes. In Studies A–C, insects initially fed on the nodules of clinically affected donor cattle. In Studies D and E, insects were initially fed via an artificial feeding membrane system (AFMS) using blood spiked with LSDV. In Study A, *Ae. aegypti* mosquitoes or *S. calcitrans* flies were interrupted during feeding on donors and transferred to naïve recipients to feed to repletion. In Study B, *Ae. aegypti* mosquitoes or *S. calcitrans* flies were fed to repletion on donors, incubated for 4 days, and then fed on recipients. Study C used only *S. calcitrans* with a 2-day incubation between donor and recipient feeding. In Study D, *Ae. aegypti* fed on an artificial membrane feeding system (AFMS), were interrupted and transferred to recipients. In Study E, *Culicoides nubeculosus* were fed via AFMS, incubated for 4 days, then fed on recipients. Animals that developed skin nodules were defined as clinical affected and represented in the figure in tan. Subclinically affected animals (blue) had LSDV DNA detected in venous blood and/or skin biopsies in the absence of any skin nodules. Non-clinical animals did not develop skin lesions and did not have detectable LSDV DNA in blood or skin (no colour).

**Table 1.**
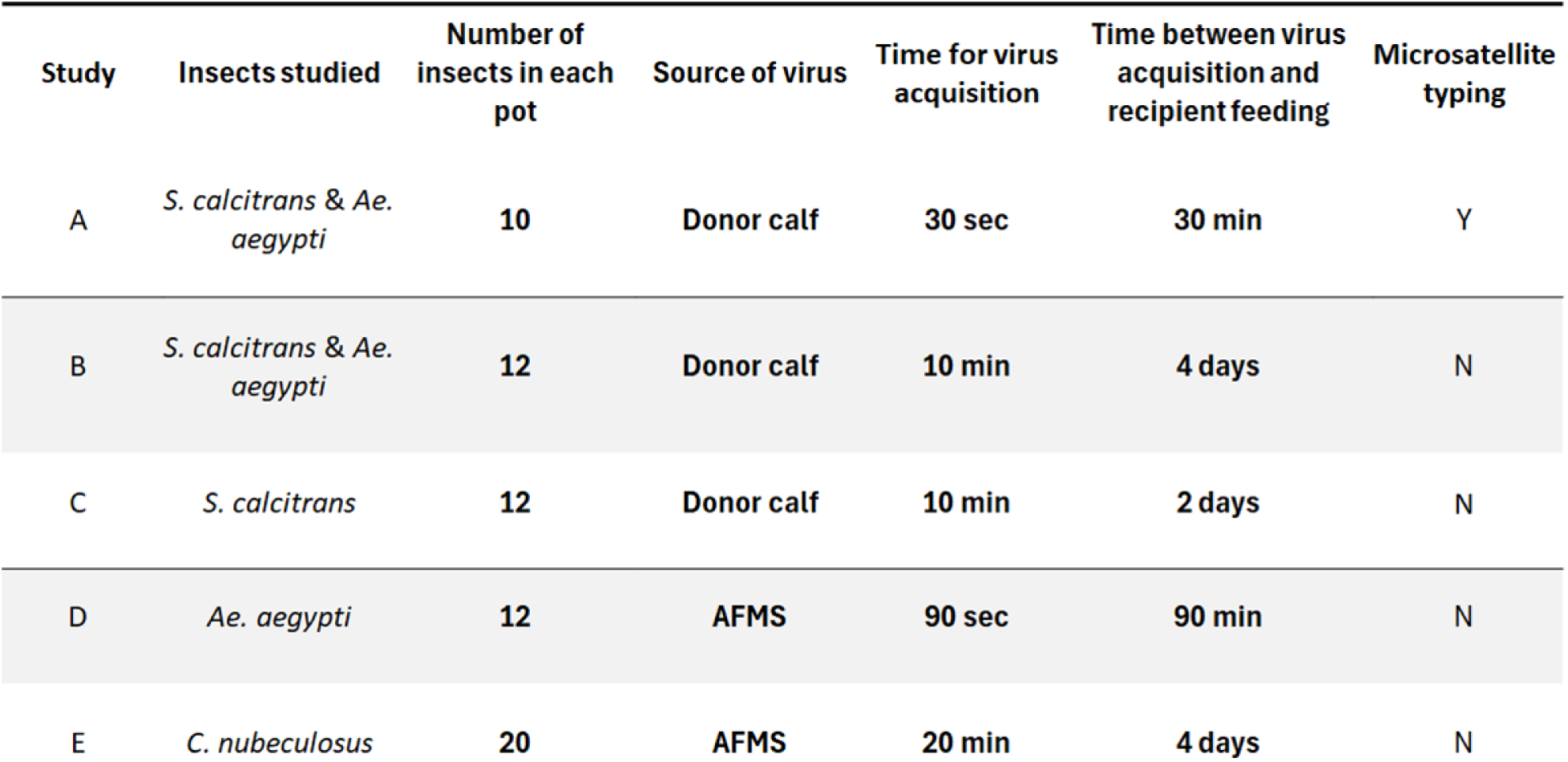
Summary of the animal studies.

| Study | Insects studied | Number of insects in each pot | Source of virus | Time for virus acquisition | Time between virus acquisition and recipient feeding | Microsatellite typing |
| --- | --- | --- | --- | --- | --- | --- |
| A | *S. calcitrans* & *Ae. aegypti* | 10 | Donor calf | 30 sec | 30 min | Y |
| B | *S. calcitrans* & *Ae. aegypti* | 12 | Donor calf | 10 min | 4 days | N |
| C | *S. calcitrans* | 12 | Donor calf | 10 min | 2 days | N |
| D | *Ae. aegypti* | 12 | AFMS | 90 sec | 90 min | N |
| E | *C. nubeculosus* | 20 | AFMS | 20 min | 4 days | N |

### Transmission of LSDV by S. calcitrans

Transmission of LSDV to a naïve recipient was studied for *S. calcitrans* after different feeding intervals: 30 min, 2 d and 4 d. Study A examined the immediate transmission of LSDV from donor to recipient calf by *S. calcitrans*. Clinically-affected donor calves were generated by needle-inoculation with LSDV as described in the materials and methods. Ten laboratory-reared *S. calcitrans* flies in a clear-walled, gauze-covered plastic pot were fed on cutaneous nodules present on the skin of a clinical donor animal for 30 sec. The pot was then removed from the nodule to “interrupt” the feed. The pot of partially fed flies was transferred immediately (within 30 min) to the skin of a recipient calf and the flies allowed to refeed for 10 min. The pot was then taken to the laboratory and stored at -80°C until analysis. Each recipient calf had two pots (20 flies) placed on either side of the dorsal midline, each day for a total of five consecutive days, giving a total of 100 flies placed on each of the five recipient calves. The time between donor and recipient feeding (30 min) mimics spread of LSDV a short distance, for example within a herd or from an infected to a neighbouring herd.

Four of the five recipient calves developed clinical LSD 11-12 days post infection (DPI) (**Figure 2**) accompanied by a viraemia and humoral immune response (**Figure 3**). Severe disease (>100 cutaneous lesions per animal) was evident in calves RS1, RS2 and RS4, and mild disease in RS5 (1 lesion). Lesions were first noted 12 (RS1, RS2 and RS4) or 13 (RS5) DPI. This indicated that *S. calcitrans* are able to transmit LSDV from a donor calf to a naïve recipient, consistent with published studies (Sohier et al., 2019, Issimov et al., 2020). No cutaneous lesions were detected on the nonclinical calf (RS3) and this calf remained viraemia negative throughout the study. Low amounts of LSDV genome were detected in a biopsy of normal skin from RS3 at 15 DPI. The amount detected was over 6log_10_ less than that detected in biopsies of cutaneous nodules on calves RS1, RS2, RS4 and RS5 (**Figure 3C**). A rising titre of neutralising antibodies was present in serum collected from calf RS3 from 7 DPI, indicating an immune response to the virus.

**Figure 2.**
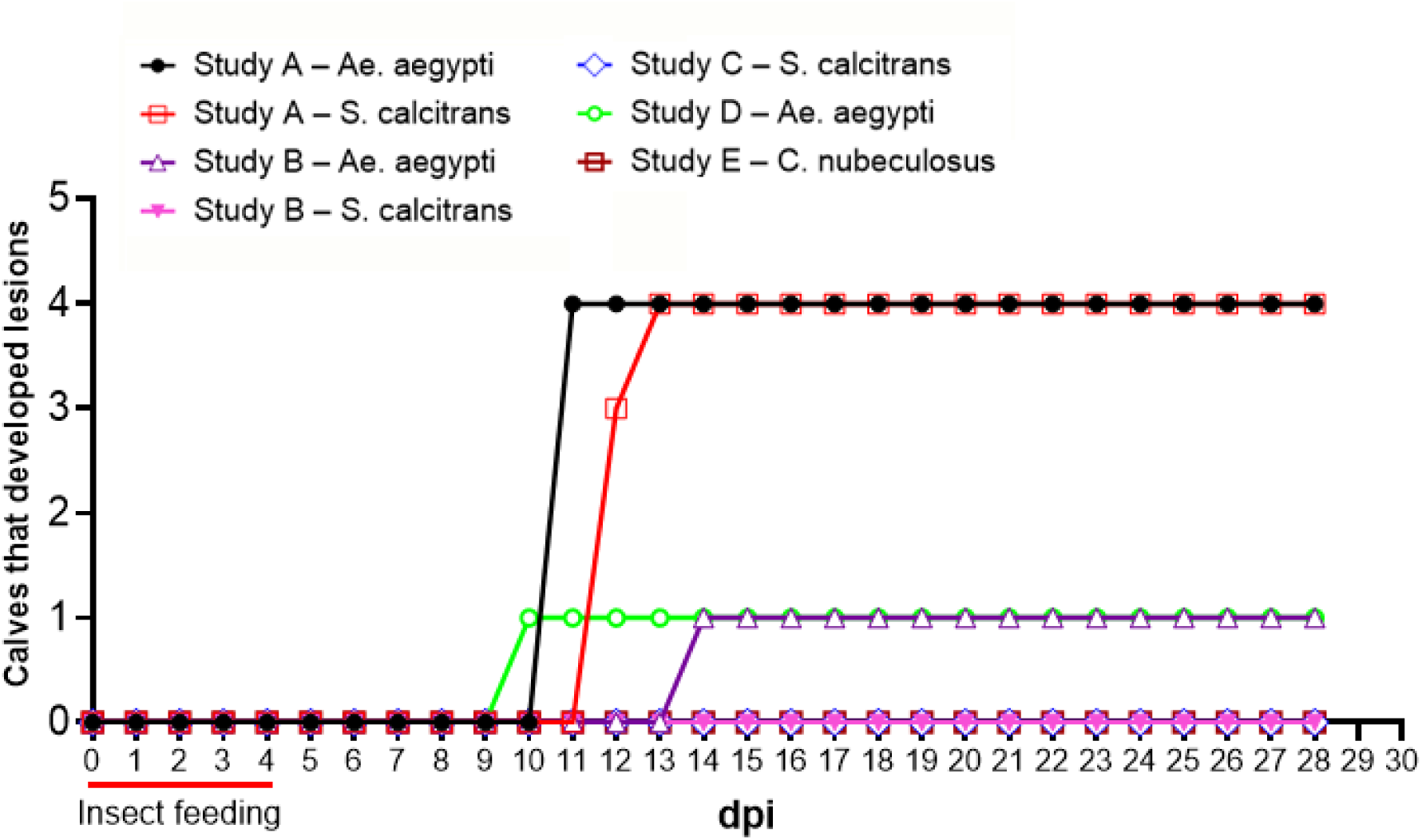
The length of time between insect feeding and development of cutaneous nodules (incubation period) in the recipient calves from all five studies (A-E). No recipient calves developed cutaneous nodules in study B (*S. calcitrans* group), study C, or study E. Six recipient calves in study A were euthanised prior to the d28 time point – RA1 (d21), RA3 (d21) and RA4 (d25 from the *Ae. aegypti* group and RS1 (d25), RS2 (d25) and RS4 (d21) from the *S. calcitrans* group.

**Figure 3.**
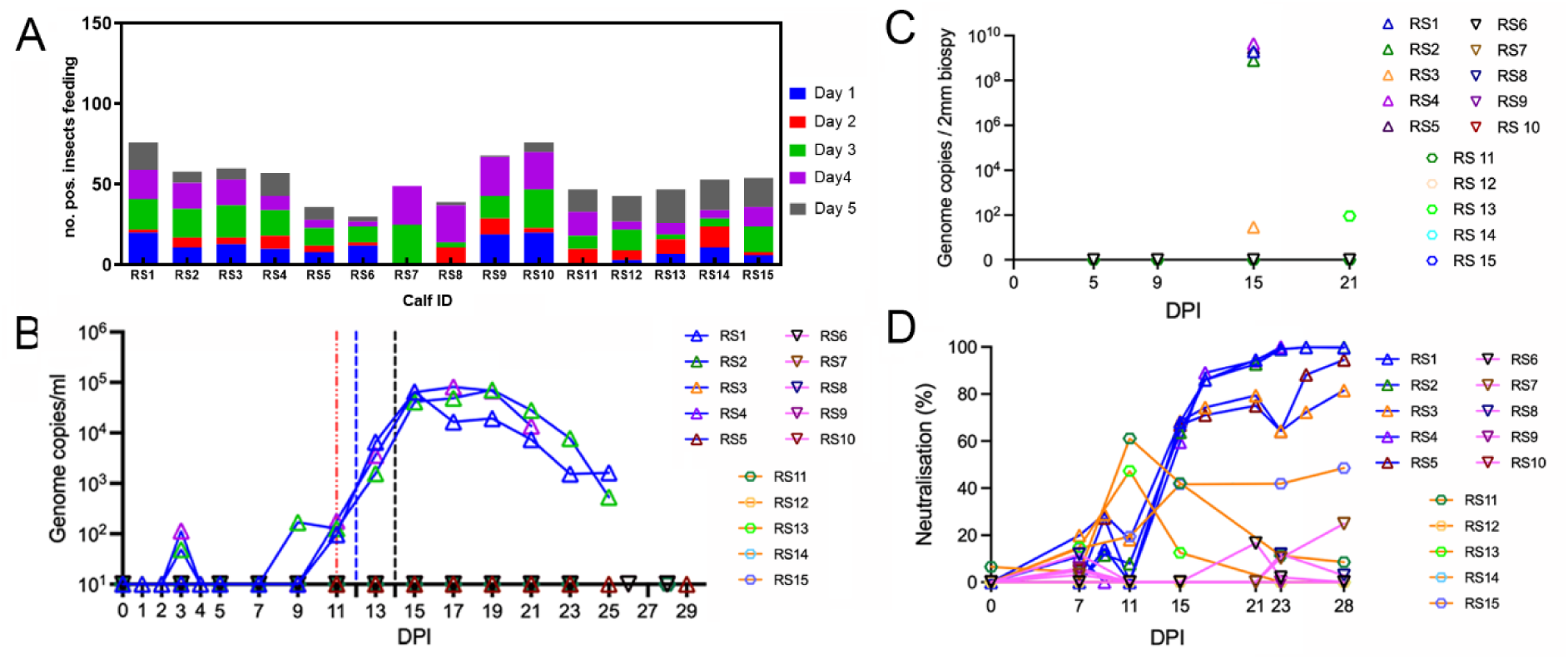
(A) The number of virus positive *S. calcitrans* that fed on each recipient calf on each of the feeding days in study A (calves RS1-RS5), study B (calves RS6-RS10) and study C (calves RS11-RS15). (B and C) Detection and quantification of LSDV DNA genome copies in the blood and skin of calves was determined by qPCR. In study A multiple punch biopsies taken on 15 DPI. The biopsy with the highest amount of virus genome detected is depicted in the figure. In studies B and C single punch biopsy samples were taken for virus genome detection on 5, 9, 15 and 21 DPI. The dotted vertical lines in (B) indicate the time points when nodules were first detected in the clinical calves. (D) Detection of partially neutralising antibodies in the sera of the recipient calves. The ability to completely or partially neutralise a fluorescently-tagged LSDV virus was carried out on MDBK cells using heat inactivated sera collected from recipient calves.

The calf which developed mild clinical disease with only a single cutaneous lesion (RS5) remained viraemia negative throughout the study across all 17 time points where blood was sampled (**Figure 3B**). The one nodule detected on the skin was biopsied on day 15 and contained high levels of LSDV genomic DNA, confirming the pathology was caused by LSDV (**Figure 3C**). This calf developed neutralising antibodies to LSDV at a lower magnitude compared to the three more severely affected calves, but a higher magnitude than the nonclinical calf RS3 on 25 and 28 DPI (**Figure 3D**).

The 500 flies (100 from each recipient calf) were analysed for presence and quantity of LSDV DNA using a qPCR assay. The flies which had acquired LSDV were then further analysed for bovine microsatellite markers to determine whether the fly had fed only on the donor calf or also on the recipient calf. This enabled accurate quantification of the number of virus-positive flies that had fed on the recipient calves.

A total of 287 out of the 500 flies had acquired virus from the donor nodule during the initial 30 sec feed, indicating very efficient acquisition of the virus. By combining the qPCR result and the microsatellite typing of the bovine blood present in each fly, we calculated that between 36 and 76 flies acquired LSDV from the donor and subsequently fed on one of the recipient calves (**Figure 3A**, calves RS1-RS5). There was no correlation between the number of virus-positive flies that fed on each recipient calf and clinical outcome, timing, or intensity of clinical disease.

To estimate the length of time LSDV can be transmitted by *S. calcitrans* flies after feeding on the donor the study was repeated (Study B) following a similar design, this time including a four-day interval between placing flies on the donor and recipient animal. The outcome of this study was very different with none of the five recipient calves developing clinical disease. The experiment was then undertaken a third time (Study C), this time with a two-day interval between placing the flies on the donor and recipient animal. Again, none of the five recipient calves developed clinical disease (**Figure 1**).

The number of virus-positive flies from studies B and C was determined using a LSDV specific PCR, indicating how many flies had acquired LSDV from the donor calf. LSDV DNA was detected in 254 flies (out of 1200; 21%), indicating efficient acquisition of virus. The presence of bovine blood in the flies was detected using a qPCR for bovine cytochrome C, used as an indicator that the flies had fed on the recipient calves. These analyses showed that each recipient calf had been fed on by between 30 and 54 virus-positive flies (Figure 3A, RS6-RS15). These results confirm that *S. calcitrans* retains LSDV genomic material for at least 4 DPI after feeding on a cutaneous nodule, and that LSDV-positive *S. calcitrans* will refeed on a naïve animal after 2 or 4 DPI. However, this did not result in the development of clinical disease in any of the ten recipient calves.

Viraemia was not detected in any of the ten *S. calcitrans* recipient calves in study B and C (**Figure 3B**). Analysis of 2mm skin biopsies taken from normal skin at days 5, 9, 15 and 21 DPI revealed the presence of LSDV DNA in only one sample (calf RS13 at 21 DPI) (**Figure 3C**).

Neutralising antibodies were detected in five of the ten *S. calcitrans* recipient calves from studies B and C (RS6, RS7, RS11, RS13 and RS15) (**Figure 3D**) indicating stimulation of a humoral immune response to LSDV. These studies indicate that biting flies are capable of transmitting LSDV soon after they acquire the virus (study A), but the likelihood of transmission is much lower two days or more post-acquisition (study B and C).

### Transmission of LSDV by *Aedes aegypti*

Study A also examined the ability of *Ae. aegypti* to immediately transmit LSDV to recipient calves. On five consecutive days ten mosquitoes in a clear-walled, gauze covered pot were fed on a cutaneous nodule on a donor calf for 30 sec. The pot was then removed from the nodule, interrupting the feed, and the partially fed mosquitoes were transferred to feed on a recipient calf within 30 min. The pot of mosquitoes was held on the recipient calf for 10 min. The pot was then stored at -80°C until analysis. The feeding process was repeated each day for a total of five consecutive days. Two pots of 10 mosquitoes were fed on each recipient calf each day, giving a total of 100 mosquitoes that had been placed on a donor nodule then on a recipient calf. Four of the five recipient calves developed cutaneous nodules at 11 DPI accompanied by a viraemia and humoral immune responses. Severe disease (>100 cutaneous lesions per animal) was evident in calves RA1, RA3, and RA5 and moderate disease in calf RA4 (50 lesions). Lesions were first noted on 11 DPI on the four clinical animals (RA1, RA3, RA4 and RA5). This is consistent with a previous study (Chihota et al., 2001) which showed that that *Ae. aegypti* were able to transmit LSDV from a donor calf to a naïve recipient. Cutaneous biopsies were taken from nodules of clinical calves RA1, RA3, RA4 and RA5 on 15 DPI and abundant amounts of LSDV genome were detected (**Figure 4C**). No cutaneous lesions were detected on the nonclinical calf (RA2), and this calf remained viraemia negative throughout the study. A cutaneous biopsy of normal skin was taken on 15 DPI and no LSDV genome was detected (**Figure 4C**). A neutralising antibody response in the nonclinical calf (RA2) was detectable from 5 DPI, but of lower magnitude compared to the clinical animals (**Figure 4D**). This is similar to the antibody response seen in the *S. calcitrans-*fed nonclinical calf RS3.

**Figure 4.**
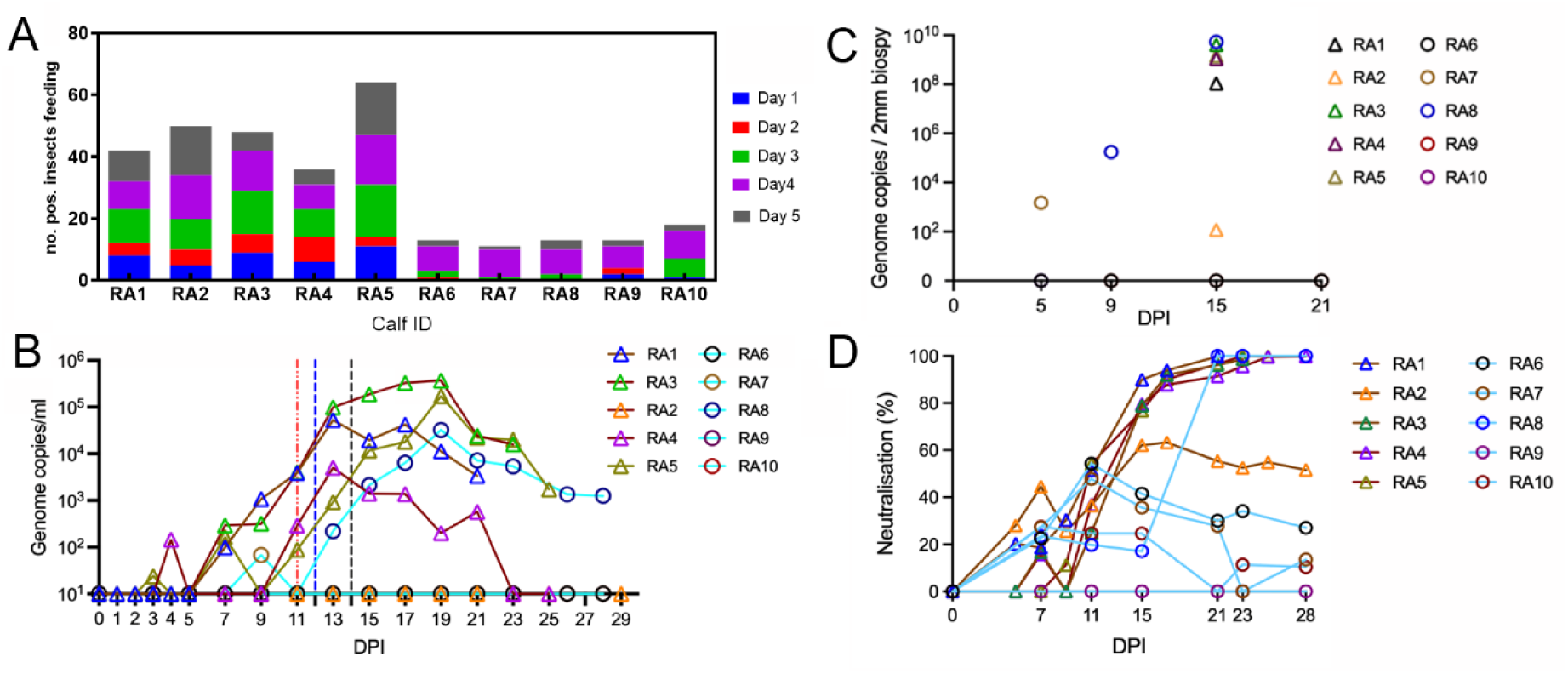
(A) The number of virus positive *Ae. aegypti* that fed on each recipient calf on each of the feeding days in study A (calves RA1-RA5) and study B (calves RA6-RA10). (B and C) Detection and quantification of LSDV DNA genome copies in the blood and skin of calves was determined by qPCR. In study A multiple punch biopsies taken on 15 DPI. The biopsy with the highest amount of virus genome detected is depicted in the figure. In study B single punch biopsy samples were taken for virus genome detection on 5, 9, 15 and 21 DPI. The dotted vertical lines in (B) indicate the first day that skin nodules were detected (11 and 12 DPI in study A, and 14 DPI for calf RA8 in study B). (D) Detection of partially neutralising antibodies in the sera of the recipient calves. The ability to completely or partially neutralise a fluorescently-tagged LSDV virus was carried out on MDBK cells using heat inactivated sera collected from recipient calves.

The 500 mosquitoes (100 from each of the five recipient calves) were analysed individually for the presence of LSDV genomic DNA as described previously (Sanz-Bernardo et al., 2021). A total of 240 mosquitoes (48%) had LSDV genomic DNA present, indicating very efficient acquisition of the virus from the nodule during the 30 sec feed. The genome copy number (**supplementary data**) indicated that mosquitoes acquired less virus in comparison to flies, possibly associated with the larger blood meal taken by flies. The mosquitoes which had acquired LSDV were analysed for bovine microsatellite markers as described above to enable accurate quantification of the number of virus-positive mosquitoes that had fed on the recipient calves. For each individual recipient calf between 36 and 64 mosquitoes had acquired LSDV and fed on each of the recipient calves (**Figure 4A, calves RA1-RA5**). There was no correlation between the number of virus-positive mosquitoes that fed on each recipient calf and the clinical outcome. For example calf RA2 was the only calf that did not develop clinical disease but had the second highest number of successful mosquito feeds (50). Similarly to the calves fed on by *S. calcitrans*, there was no difference in the timing or intensity of clinical disease that correlated with the number of virus-positive mosquitoes that fed on each recipient calf. The parameters of clinical disease in the four clinical calves were compared with the clinical disease in the four clinical calves fed on immediately by virus-positive *S. calcitrans* and no difference was detected, indicating that feeding of virus-positive mosquitoes and flies leads to indistinguishable disease in recipient calves.

The experiment was repeated in Study B, this time incorporating a four-day interval between feeding of *Ae. aegypti* on donor and recipient calf. Twelve mosquitoes were placed in each pot, then fed on a cutaneous nodule on a donor animal for 10 min to enable feeding to repletion. The pots of mosquitoes were then incubated for 4 days before being fed on naïve recipient calves, again for 10 min. Two pots of mosquitos were fed on each recipient calf every day for five sequential days, a total of 120 mosquitoes per recipient calf. In contrast to study A where four recipient calves developed clinical disease, only one of the five recipient calves developed clinical disease in study B, calf RA8. This result showed that *Ae. aegypti* mosquitoes may transmit LSDV for at least 4 days following virus acquisition. The incubation period of disease in RA8 was longer than seen in the recipients fed on by virus-positive *Ae. aegypti* in Study A, with cutaneous nodules first identified at 14 DPI rather than 11 DPI (**Figure 2**). The disease was also milder than that seen in study A, with only 20 nodules detected compared with over 100 nodules on the majority of recipient calves in study A. Viraemia in calf RA8 was also detected later at 13 DPI and was briefer with a lower magnitude compared to the four clinical calves in study A (**Figure 4B**). The only other calf with a detectable viraemia in study B was RA10, with approximately 10^2^ genome copies detected at 9 DPI.

Skin biopsies (2mm) were taken from each recipient calf on 5, 9, 15 and 21 DPI and tested for the presence of LSDV DNA. At 5 DPI a biopsy was taken from normal skin of each calf. LSDV DNA was detected in one sample from calf RA10 (which subsequently developed LSDV viraemia). At 9, 15 and 21 DPI biopsies were taken from cutaneous nodules of clinical calf RA8 and abundant LSDV DNA was detected (**Figure 4C**). No LSDV DNA was detected in the biopsies of normal skin taken on 9, 15 and 21 DPI from the four nonclinical calves.

Each mosquito from study B was analysed using PCR for LSDV genomic DNA and bovine cytochrome b DNA to determine the number of virus-positive mosquitoes that had fed on the recipient calf. Fewer mosquitoes were virus-positive after an incubation of 4 days, consistent with our previous findings (Sanz-Bernardo et al., 2022, Sanz-Bernardo et al., 2021) with 68 out of 600 mosquitoes virus-positive after recipient feeding. A similar number of virus-positive mosquitoes fed on each calf, varying from 11 mosquitoes on RA7 to 18 mosquitoes on RA10 (**Figure 4A, calves RA6-RA10**). Feeding was most successful on day 4 with 60% of virus-positive mosquitoes feeding on recipients, and least successful on day 1 with only 4.4% of virus-positive mosquitoes feeding on recipients. A total of 13 virus-positive mosquitoes had fed on RA8, indicating only a small number of virus-positive insects are needed to cause disease.

We quantified antibodies to LSDV in the serum of all five recipient calves in study B using a fluorescent virus neutralisation test (Fay et al., 2022) (**Figure 4D**). As expected, a rising titre of neutralising antibodies over time was detected in the serum of the clinical calf RA8 with total neutralisation of input virus detected by 21 DPI. Lower levels of neutralising antibodies were detected in three of the other calves (RA6, RA7 and RA10) from 7 DPI, peaking at 11 DPI. No neutralising antibodies were detected at any time point in the serum of calf RA9.

These results indicate that calves RA6, RA7 and RA10 were exposed to LSDV from the virus-positive mosquitoes but did not go on to develop clinical disease or viraemia Calf RA9, despite being bitten by a similar number of virus-positive mosquitoes, did not develop an antibody response to LSDV.

Previous work has shown that the acquisition and retention of LSDV by mosquitoes fed on an artificial membrane feeding system (AMFS) containing horse blood spiked with LSDV was similar to mosquitoes fed on nodules on clinical calves (Sanz-Bernardo et al., 2022). In Study D we examined whether mosquitoes that had acquired LSDV from an AMFS could cause LSD in recipient calves. The study design was similar to Study A with the AMFS replacing donor calves as the source of virus. Mosquitoes were fed on the AMFS for 90 sec then feeding was interrupted, and the mosquitoes transferred as soon as possible (within 90 min) to the animal facility to feed on five recipient calves. Twenty-four mosquitoes (12 mosquitoes in two pots) were fed on each recipient calf each day for 5 consecutive days, giving a total of 120 mosquitoes per recipient calf. After each feeding session the mosquitoes were stored at -80°C until analysis.

Analysis of the mosquitoes revealed between 8 and 14 mosquitoes had acquired LSDV from the AMFS and then fed on the recipient calf (**Figure 5A**). One of the five recipient calves (RAV4) developed clinical disease with cutaneous nodules first detected at 10 DPI (**Figure 2**). Only 14 mosquitoes had acquired LSDV and fed on RAV4, indicating again that very few virus-positive mosquitoes are required to cause LSD. The clinical disease exhibited by RAV4 was indistinguishable from that seen in recipient calves infected by mosquitoes in Study A, with lymphadenopathy and pyrexia preceding the appearance of cutaneous nodules. Calf RAV4 had detectable levels of LSDV DNA in the blood stream from 9 DPI (**Figure 5B**). No LSDV DNA was detected in the biopsies of normal skin taken from calf RAV4 at 5 and 9 DPI, prior to the development of lesions, however abundant LSDV DNA (over 10^10^ genome copies per biopsy) was present in the biopsy of cutaneous nodules at 15 and 21 DPI (**Figure 5C**). No viraemia was detected in any of the other four calves at any point in the study (**Figure 5B**).

**Figure 5.**
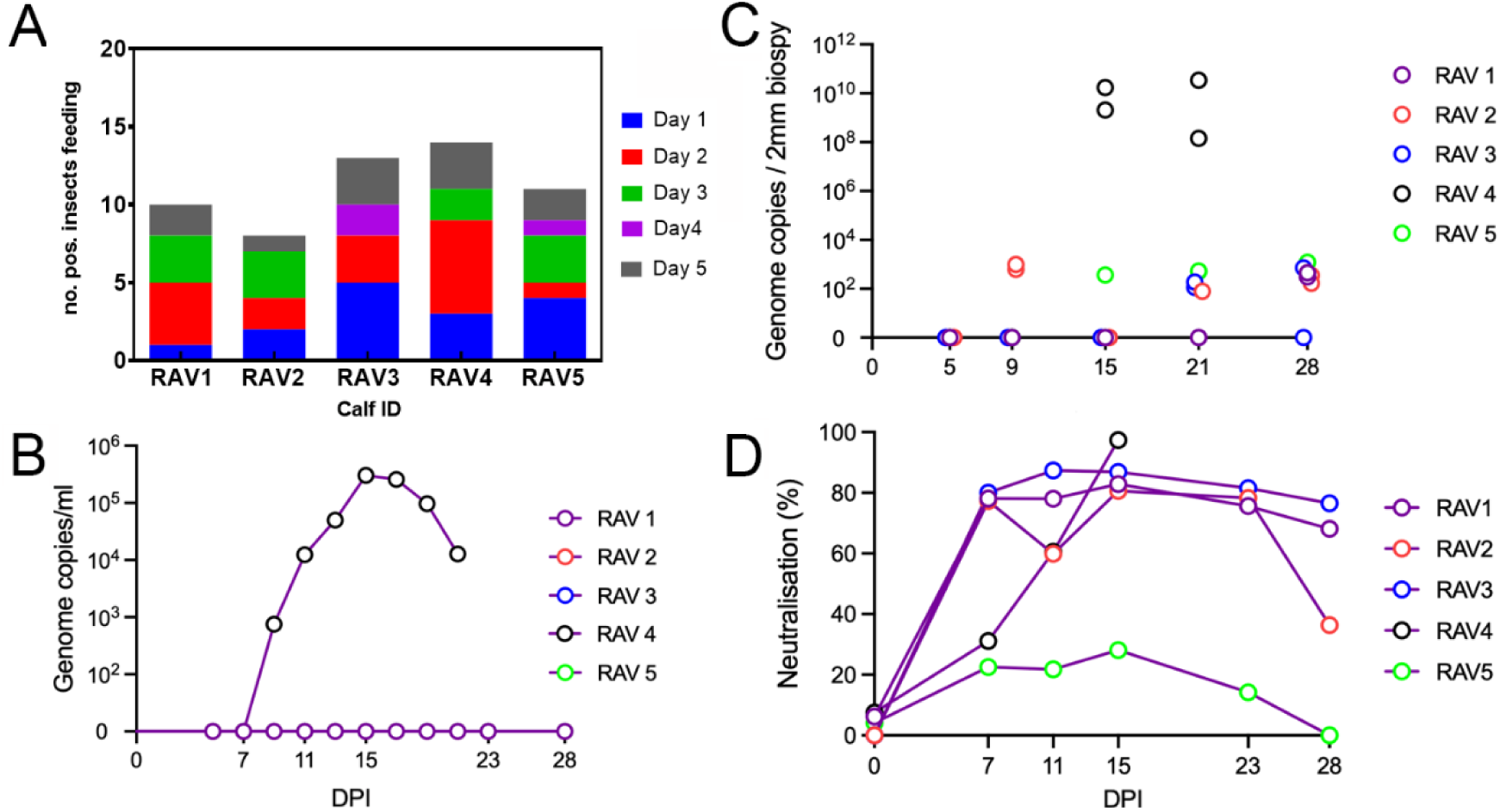
(A) The number of LSDV-positive *Ae. aegypti* that fed on each of the five recipient calves (RAV1-5) on each of the feeding days. The *Ae. aegypti* were fed daily on LSDV spiked horse blood via an artificial membrane feeding system for 90 seconds. The mosquitoes were immediately transferred to feed on naïve recipient calves (RAV1-5). This was repeated each day for a total of 5 days. (B and C) Detection and quantification of LSDV DNA genome copies in the blood and skin of calves was determined by qPCR. The dotted vertical line indicates the first day that skin nodules were detected on calf RAV4. In (C) the data points for RAV2 and RAV3 have been jittered to allow visibility. (D) Detection of partially neutralising antibodies in the sera of the recipient calves. The ability to completely or partially neutralise a fluorescently-tagged LSDV virus was carried out on MDBK cells using heat inactivated sera collected from recipient calves.

However LSDV DNA was detected in normal skin tissue from all four non-clinical calves during the study duration. Samples of normal skin from calf RAV2 and RAV5 were positive at 9, 15, 21 and 28 DPI, RAV3 was positive on 21 DPI, and RAV1 positive on 28 DPI (**Figure 5C**). Only low amounts of virus were detectable in these biopsy samples, between 10^1^ and 10^3^ genome copies per biopsy. High levels of neutralising antibody were present in the serum of the calf RAV4 that developed clinical disease (calf RA4, **Figure 5D**). Consistent with the skin biopsy results, all four non-clinical calves developed relatively high titres (around 80% of input virus neutralised) of neutralising antibodies to LSDV during the study.

### Transmission of LSDV by *Culicoides nubeculosus*

The final study, study E, examined the ability of the midge species *C. nubeculosus* to transmit LSDV. Our previous work indicated that LSDV could be acquired by midges from the nodules of clinical calves and retained for up to four days (Sanz-Bernardo et al., 2021). However, no studies have reported successful transmission of LSDV by any midge species. The life cycle of midges means they do not refeed until oviposition has taken place, around 4 days after an initial feed, depending on the ambient temperature. The study was therefore designed with a four-day interval between first and second feed. At least 15000 midges were fed on an AMFS spiked with LSDV each day for five days. Immediately after the first feed midges were rendered immobile by CO_2_ and those that had fed were sorted by eye. On each day 500 blood-fed midges were sorted into 5 pots (100 midges per pot) and the pots incubated at room temperature for 4 days. On the fourth day the midges were transported to the animal facility, placed on naïve recipient calves and allowed to feed for 20 min. After feeding the midges were stored at -80 °C until analysis.

Analysis of the midges revealed that 62 midges were virus-positive, confirming previous work which showed that midges remain positive for LSDV DNA for at least four days post feeding. Between 1 and 20 virus-positive midges fed on each of the recipient calves (**Figure 6A**). None of the five recipient calves developed clinical disease (no cutaneous nodules developed and no lymphadenopathy or temperature spike was detected), and no LSDV DNA was detected in the bloodstream or in any skin biopsy sample taken from any calf across the duration of the experiment. Immunological analysis revealed neutralising antibodies capable of neutralising at least 50% of the input virus in two animals (RM1 and RM4). Lower titres (neutralising between 20-40% of input virus) were detected in the remaining three calves, with highest titres 23 and 28 DPI (**Figure 6B**).

**Figure 6.**
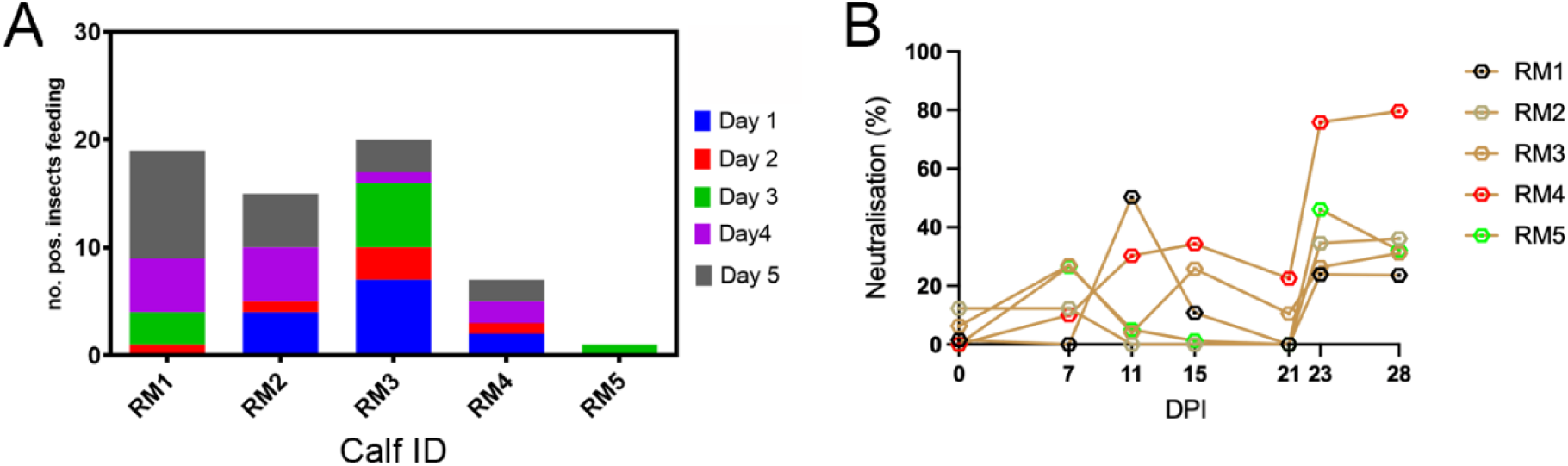
(A) The number of LSDV-positive *C. nubeculosus* that fed on each of the five recipient calve (RM1-5) on each of the feeding days. The *C. nubeculosus* were fed daily on LSDV spiked horse blood via an artificial membrane feeding system for 20 minutes. The midges were then incubated for 4 days then fed on naïve recipient calves (RM1-5). This sequence was repeat each day for a total of 5 days. (B) Detection of partially neutralising antibodies in the sera of the recipient calves. The ability to completely or partially neutralise a fluorescently-tagged LSDV virus was carried out on MDBK cells using heat inactivated sera collected from recipient calves.

### Modelling of LSD transmission

For studies A-E each insect was examined individually for acquisition of LSDV and for feeding on the recipient calf. This data was then modelled, building on our previous work (Gubbins, 2019, Sanz-Bernardo et al., 2021). The model for each species yielded an adequate fit to the data (posterior predictive *P*-values >0.25 for all recipients). The posterior estimates for the probability of transmission from insect to bovine for each species are shown in Table 2. This probability was higher for *Ae. aegypti* (posterior median: 0.029) compared with *S. calcitrans* (posterior median: 0.005), indicating that a virus-positive *Ae. aegypti* is nearly 6 times more likely to transmit LSDV than a virus-positive *S. calcitrans*. The probability of transmission from *Ae. aegypti* that had fed on a donor calf and *Ae. aegypti* that had fed on an AMFS containing LSDV was similar (posterior medians: 0.029 and 0.033, respectively) (Table 2), supporting the use of AMFS to study vector transmission of LSDV. The estimate for probability of transmission from *C. nubeculosus* to cattle is zero (posterior mode), since no transmission occurred after feeding though all recipients were fed on by at least one LSDV-positive insect, however the upper 95% credible limit is 0.05 (Table 2), indicating a moderate level of uncertainty.

**Table 2.** Parameters for transmission of lumpy skin disease virus by three species of biting insect.

|  | posterior<br>median | 95% credible interval |
| --- | --- | --- |
| probability of transmission from insect to bovine (*b*) |  |  |
| *Aedes aegypti* (fed on calves) | $2.9 \times 10^{-2}$ | $(1.1 \times 10^{-2}, 6.2 \times 10^{-2})$ |
| *Aedes aegypti* (membrane-fed) | $3.3 \times 10^{-2}$ | $(5.0 \times 10^{-3}, 1.1 \times 10^{-1})$ |
| *Culicoides nubeculosus* | 0† | $4.6 \times 10^{-2} \dagger$ |
| *Stomoxys calcitrans* | $5.1 \times 10^{-3}$ | $(1.6 \times 10^{-3}, 1.2 \times 10^{-2})$ |
| basic reproduction number ($R_0$) | | |
| *Aedes aegypti* | 0.58 | (0.12, 1.20) |
| *Culicoides nubeculosus* | 0† | 7.3† |
| *Stomoxys calcitrans* | 5.8 | (0.8, 17.0) |

† posterior mode and upper 95% credible limit

Once the estimates for the probability of transmission from insect to bovine for each species had been calculated they were incorporated in calculations for the basic reproduction number (*R*_0_) (**Figure 7**; **Table 2**). This value takes into account life history parameters of each vector species including frequency of biting, vector to host ratio, and vector mortality rate. The *R*_0_ was highest for *S. calcitrans* (median 5.8) suggesting this is an important vector of LSDV. The estimated *R*_0_ for *Ae. aegypti* was below one (median 0.6) suggesting this may not be an important vector on its own but may still contribute to transmission. For *C. nubeculosus* the estimated *R*_0_ is zero (posterior mode), but the upper 95% credible limit is 7.3 (so >1), meaning uncertainty remains about whether this species could be an important vector of LSDV.

**Figure 7.**
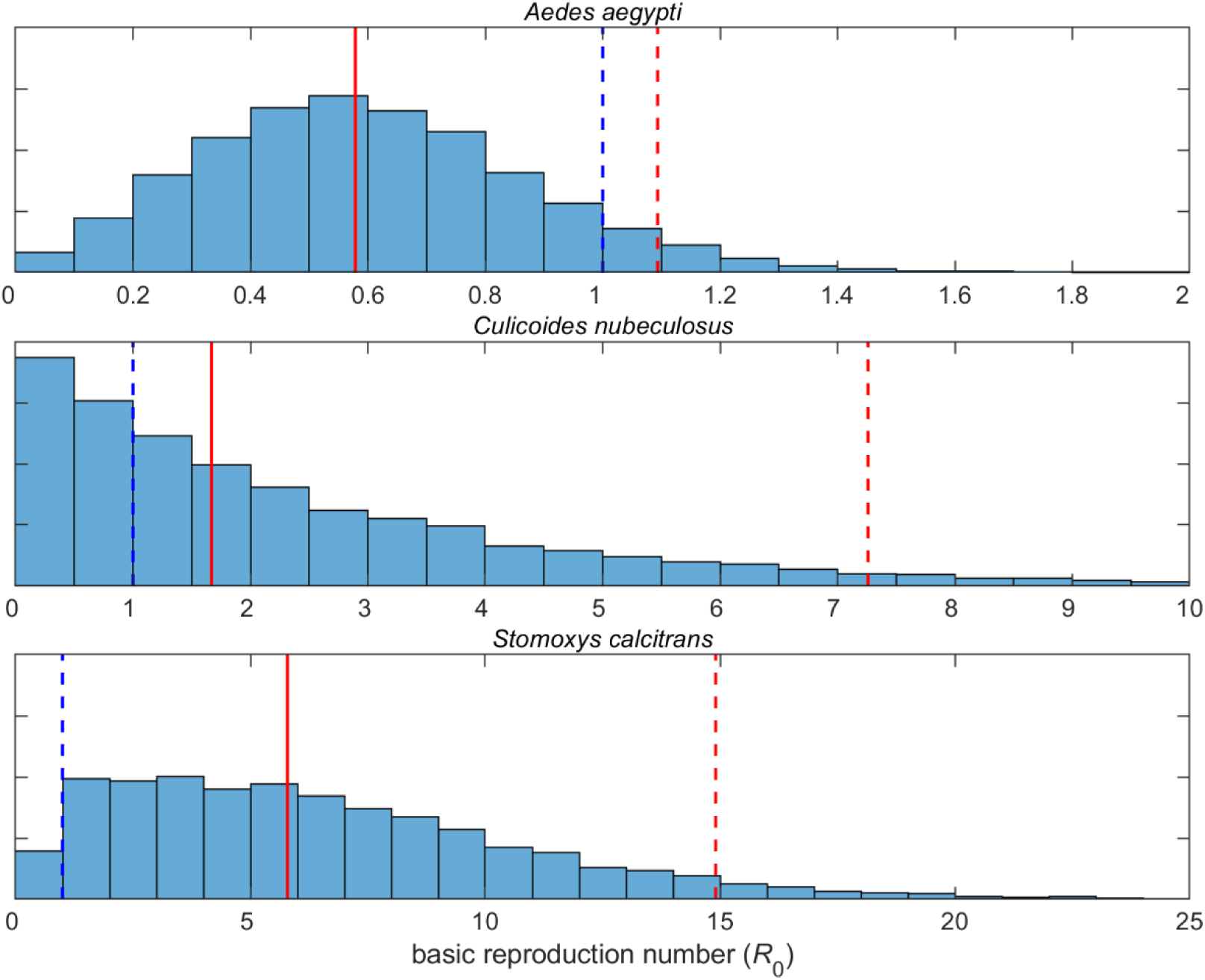
Basic reproduction number (*R*_0_) for lumpy skin disease virus when transmitted between calves by *Aedes aegypti* (top), *Culicoides nubeculosus* (middle) or *Stomoxys calcitrans* (bottom). The plot shows a histogram for *R*_0_ based on replicated Latin hypercube sampling (100 replicates with the range for each parameter subdivided into 100 steps). The solid and dashed red lines show the median and upper 95% credible limit, respectively. The blue dashed line indicates the threshold at *R*_0_=1.

## Discussion

An incomplete understanding of the vector transmission of LSDV is one of the most important knowledge gaps preventing control of this emerging transboundary disease. In this work we have shown that *Ae. aegypti* and *S. calcitrans* are capable of transmitting the virus and causing disease in a recipient animal under experimental conditions. *C. nubeculosus* midges did not transmit disease but did produce a LSDV-specific immune response. Only individual insects that take a blood-meal both first on the donor and secondly on the recipient can in principle transmit LSDV mechanically. Combining the detection of bovine genetic markers with the detection of LSDV we were able to quantify the number of individual insects fulfilling this feeding requirement. This individual analysis of insects post-feeding enabled calculations of the likely ability of each vector species to transmit the virus in the field. Our results validate that *S. calcitrans* and, by extension, other biting flies are likely to be important vectors for transmitting LSDV.

Our initial experiment (study A) mimicked short-distance transmission by *S. calcitrans* and *Ae. aegypti*. Similar numbers of *S. calcitrans* and *Ae. aegypti* fed on donor calves, acquired LSDV and transmitted it to naïve recipient animals in study A. Disease outcomes were similar between the *S. calcitrans* and *Ae. aegypti* recipient groups, with no indication that the species of vector influences subsequent pathology. There was also no correlation between the numbers of virus-positive insects that fed on the recipient and their subsequent clinical status, suggesting that the factors which determine whether or not a calf develops LSD after blood-feeding by a virus-positive insect are complex, involving more than simply the number of virus-positive insects which have fed on the calf. For example it is possible that LSDV present on the mouthparts of the insect may be the only virus transmitted to a recipient during biting, with LSDV present elsewhere in the insect, such as in the abdomen, unable to reach the skin of the recipient and therefore not playing a role in transmission. This would result in “virus-positive” insects, representing insects with LSDV present in any part of their anatomy, over-representing insects able to transmit virus. *C. nubeculosus* was not included in this initial short-distance transmission experiment as *Culicoides* midge species delay their refeeding to follow the laying of eggs (Rozo-Lopez et al., 2022), so would be very unlikely to refeed on a recipient only 30 min after feeding on the donor.

Probably the most important outcome from study A was how efficient insects are in transmitting LSDV. LSD is notoriously difficult to replicate experimentally, with large challenge doses (in the region of 10^6^ pfu per animal) resulting in approximately 50% of animals developing clinical disease (Moller et al., 2019, Sohier et al., 2019, Sanz-Bernardo et al., 2021, Sanz-Bernardo et al., 2022). However, in study A, which used vector transmission, 80% of cattle developed clinical disease. The features of vector transmission which potentiate development of LSD are unknown but may include co-inoculation of pro-viral factors in the saliva, inoculating just the right amount of virus, or inoculating the virus into exactly the right part of the skin.

After establishing the ability of *S. calcitrans* and *Ae. aegypti* to transmit LSDV, we next asked how long these vectors remain able to do so. A set of studies (B, C and E) measuring the transmission ability of vectors after 2 or 4 days incubation were carried out with *Ae. aegypti, S. calcitrans* and *C. nubeculosus*. Only one calf out of 20 developed clinical disease. This calf had been fed on by 8 virus-positive *Ae. aegypti* mosquitoes four days after they acquired the virus. This positive finding suggests that mosquitoes may play a role in long distance spread of LSDV, with mosquitoes having an average maximum flight distance of between 50 m and 50 km depending on the species (Verdonschot and Besse-Lototskaya, 2014). In contrast, all ten calves fed on by virus-positive *S. calcitrans* and five calves fed on by virus-positive *C. nubeculosus* remained clinically normal. This ability of *Ae. aegypti* to transmit LSDV very efficiently was captured in the calculation of the probability of transmission. A virus-positive *Ae. aegypti* is nearly six times more likely to cause LSD in a recipient than a virus-positive *S. calcitrans*. It is not clear why mosquitoes are more efficient than biting flies, it is possible the different feeding mechanisms may contribute. Mosquitoes make a tiny incision into the skin and direct their proboscis into a capillary (Krenn and Aspöck, 2012), whereas biting flies generate a pool of blood and therefore greater damage to the skin.

The clinical, virological and immunological analyses shed more light on the status of the recipient calves which did not develop clinical disease. Virus transmission in the absence of clinical disease was identified in calf RS3 in study A with virus genome detected in the skin biopsy taken 15 days after *S. calcitrans* feeding, and RA10 in study B after being fed on by virus-positive mosquitoes. LSDV genome was detected in both blood and skin of this calf.

Viral DNA was also detected in one recipient in study C, calf RS13, in a skin biopsy taken at 21 days post feeding. Antibodies capable of neutralising LSDV were detected in all three of these nonclinical calves (RS3, RA10 and RS13). Neutralising antibodies were also detected in three non-clinical recipients that had been fed on by *Ae. aegypti* after a four-day incubation, one non-clinical recipient that had been fed on by *S. calcitrans* after a four day incubation, three non-clinical recipients that had been fed on by *S. calcitrans* after a two day incubation, and all five recipients fed on by *C. nubeculosus*. These recipient calves had therefore been exposed to LSDV, or components of LSDV, by the vector but did not go on to develop disease. The basis for this resistance is unknown, but is likely due to one or more factors that influence the complex virus – host – vector relationship.

The detection of viral genome in a skin biopsy from calf RS13 at just one time point late in infection (21 DPI) was unexpected as this animal had no clinical signs and no viral genome was detected in the blood at any time point. LSDV genome has previously been shown to be present only intermittently in the skin and blood of subclinically infected animals (Haegeman et al., 2023, Aerts et al., 2021, Sanz-Bernardo et al., 2021), and the moderate neutralising antibody titre detected in RS13 is consistent with subclinical infection. An incubation period of greater than 20 days has been estimated in previous LSDV vector transmission studies (Sohier et al., 2019, Haegeman et al., 2023), and together with our results this suggests that the incubation period of LSD following vector transmission could be longer than previously estimated from needle-inoculated experimental studies.

In studies A-C it was necessary to generate a group of donor calves with cutaneous nodules in order to study the vector transmission of LSDV. However previous work has shown that vectors efficiently acquire LSDV from a AMFS (Sanz-Bernardo et al., 2022), therefore we examined whether an AMFS could replace the donor calves in the experimental design (study D). One of the five recipient calves in this study developed clinical disease, providing proof of principle that this design works. However the rate of disease incidence was lower compared to study A, where four out of five calves developed clinical disease following immediate donor to recipient transmission. Analysis of the insects revealed that an average of 8-14 LSDV positive mosquitoes fed on each recipient calf after acquiring virus from the AFMS, compared to 36-76 mosquitoes in study A. It is possible that optimisation of AMFS feeding of the mosquitoes would result in a higher rate of disease in recipients.

A core aim of this project was to generate robust, quantitative data describing the vector transmission of LSD. Data from each of the five studies on the exact number of insects which acquired virus and fed on recipient calves was combined with the clinical outcomes and addressed this aim, enabling a more accurate mathematical model of LSDV transmission to be generated. Mathematical modelling further took into account factors such as feeding behaviours, numbers of individual insects feeding on an individual animal and life parameters. This produced a clear result – that *S. calcitrans* and, by inference, other biting fly species are likely to be important vectors for LSDV transmission with an *R*_0_ of 5.8.

Interestingly, even though *Ae. aegypti* have a higher probability of transmission than *S. calcitrans*, once life history parameters are taken into account R_0_ is 0.6, suggesting mosquitoes may be contributors but not drivers of LSDV transmission. This difference is principally due to differences in biting rate: *S. calcitrans* take a blood meal every 4 to 72 hours compared with *Ae. aegypti* which take a blood meal every 4 to 5 days (Gubbins, 2019).The estimated *R*_0_ for *C. nubeculosus* is zero, but the upper 95% credible limit is 7.3 (so >1), meaning uncertainty remains about whether *Culicoides* biting midges could be a relevant vector of LSDV.

This study provides new knowledge about the role of different hematophagous insects in the spread of LSDV. Experimental data alone suggests that *Ae. aegypti* is capable of transmitting LSDV in both immediate and delayed transmission scenarios whereas *S. calcitrans* is capable of transmitting LSDV in immediate transmission scenarios only. Incorporating modelling to understand transmission dynamics was a powerful tool to better understand the risk posed by these different insects, highlighting biting flies as an important vector for LSDV transmission.

## Materials and methods

### Viruses

The LSDV strain used was provided by the WOAH Capripoxvirus Laboratory at The Pirbright Institute and originated from an outbreak of LSD in Eastern Europe in 2016. LSDV was propagated on Madin-Darby bovine kidney (MDBK, ATCC CCL 22 (NBL-1) *Bos Taurus*) cells and semipurified through a 36% sucrose cushion prior to titration, as described previously (Fay et al., 2020). A pestivirus, a known contaminant of LSDV strains, was present in the inoculum. Subsequent testing identified pestivirus genome in EDTA blood samples collected from donor calves but not recipient calves.

### Animals and experimental design

Cattle were housed in the primary high-containment animal facilities (biosafety level 3) at The Pirbright Institute. The husbandry of the animals during the study was described previously (Sanz-Bernardo et al., 2020). Cattle used in the studies were Holstein-Friesen male cattle, aged between 111 and 122 day old and weighing between 93 and 162 kg. The animals were housed in groups of five within a high-containment (SAPO4) animal facility. Bedding material was provided (https://www.mayofarmsystems.co.uk/mayo-mattress-stable-mat/), light/dark cycle was 12:12 hour, temperature was held between 10°C to 24°C, and humidity 40% to 70%. Animals were given ad lib access to hay and water, with concentrate rations provided twice daily. Environmental enrichment was provided including rubber toys and a hollow ball stuffed with hay. Cattle were under veterinary care for the entire duration of the study. The nonsteroidal anti-inflammatory drug meloxicam (0.5 mg/kg of body weight) (Metacam 20-mg/ml solution; Boehringer Ingelheim) was used when required on welfare grounds.

Experiments were carried out to investigate and quantify the transmission of LSDV from clinically affected “donor” calves to naïve “recipient” calves (Studies A-C, Figure 1, Table 1). Additionally two further studies (Studies D and E, Figure 1, Table 1) were carried out to further assess efficiency of LSDV transmission from artificially membrane fed insects to recipient cattle.

In study A cattle were typed using microsatellite markers (see below) and assigned into the donor or recipient group based on this. For studies B and C the calves were stratified by weight then randomly assigned using the RAND() function in Excel to donor or recipient groups. Only recipient cattle were used in studies D and E so no randomisation was carried out.

Each study varied the infection route and interval between feeding and exposure to recipient cattle. In studies A-C, LSDV infected donor cattle were the source of LSDV infection. In study A, five donor calves were challenged with 3 x 10^6^ PFU LSDV via intravenous and intradermal route (Sanz-Bernardo et al., 2021). Calves were challenged with 2ml of inoculum (1 x 10^6^ PFU/ml) intravenously and 1ml of inoculum intradermally divided in 4 locations of 0.25ml each, 2 on the left side of the neck and 2 on the right side of the neck. In both studies B and C five donor calves were challenged with 1 x 10^6^ PFU of LSDV in ten 100µl “microdoses” on both the left and right dorsal flanks of the calf just caudal to the scapula.

It was not possible to conceal the treatments allocated to each calf from the animal care staff or those administering the treatment. Laboratory scientists were aware of the group (donor or recipient) and the treatment (species of vector) but unaware of the clinical outcome of the treatment (clinical or non-clinical). Clinical outcome measures were taken from each calf at regular intervals, including rectal temperature, heart rate, respiration rate, behaviour (alertness, feeding and responsiveness), size of the superficial pre-femoral and pre-scapular lymph nodes, and presence of oral or nasal discharge. The number of cutaneous nodules was counted up to a maximum of 100 per animal.

A subset of the donor calves in each of studies A-C developed clinical disease consistent with that seen in the field and in our previous studies (Sanz-Bernardo et al., 2020, Sanz-Bernardo et al., 2021, Sanz-Bernardo et al., 2022), characterised by multifocal necrotising cutaneous nodules.

### Microsatellite marker analysis

In study A each donor and recipient calf was genotyped prior to the study commencing using bovine microsatellite markers. Calves were assigned to donor or recipient groups based on their microsatellite type, ensuring that donor and recipient calves could be distinguished using this method. Microsatellite marker analysis was then used during the study to quantify the number of insects that fed on a donor and/or a recipient. Thereby we were able to quantify the exact number of individual insects that had taken a blood-meal on the donor calf as well as on the recipient calf and hence would be the only individuals theoretically able to transmit LSDV.

Sample preparation was carried out following the manufacturer guidelines (Bovine Genotypes Panel 3.1, Thermo Scientific). DNA was extracted from oral swabs taken from study animals. Swabs were incubated with 300 µl of digestion buffer containing proteinase K for 30 min at 50°C then vortexed for 2 min, and 100 µl taken for DNA extraction. DNA extraction was performed using the MagMAX™ CORE AgGenomic DNA Extraction Kit. Genomic DNA was eluted into 50µl elution buffer and DNA concentration measured using the QuantiFluor™ ds-DNA dye (Promega) on the Cytation (BioTek) using the Lambda DNA standard provided with the kit. PCR reactions were performed on the T100 Thermal Cycler (BioRad) using Bovine Genotypes Master Mix (F-841) and Bovine Genotypes Panel 3.1 Primer Mix (F-902) (Thermo Scientific) provided with the kit and following the manufacturer instructions. DNA electrophoresis was carried out using a 3730 DNA Analyser (AB, HITACH) and GeneMapper v6.0 (Applied Biosystems) and the GeneScan 500 LIZ Size Standards [REF: 4322682, Applied Biosystems) were used to analyze the peak of the samples. The Bovine Genotypes Panel 3.1 kit can amplify 18 STR loci. For each of the animals all loci were analyzed, and the SPS113 and SPS115 loci selected to differentiate the donor/recipient groups. Additional markers based on signal intensity included ETH10, BM2113 and MGTG4B (R). The same microsatellite marker methodology was used on DNA extracted from insects post-feeding to identify if each insect had fed on the donor only, the recipient only or both the donor and the recipient.

### Insects

Hematophagous insects used in this study were *Ae. aegypti* “Liverpool” strain, *S. calcitrans* (colony established in 2011 from individuals kindly provided by the Mosquito and Fly Research Unit, USDA Florida), and *C. nubeculosus* (Boorman, 1974). Insects were reared as described previously (Sanz-Bernardo et al., 2021). *Aedes aegypti* and *S. calcitrans* were maintained at 26°C, 50% RH, and 12:12 light/dark cycle and *C. nubeculosus* were maintained at 27 ± 2°C, 50% RH, with a 16:8 light/dark cycle. The insects were supplied and used 5-7 days post eclosion for *Ae. aegypti*, 2-4 days for *S. calcitrans* and 0-2 days for *C. nubeculosus*.

### Insect feeding

All insects were starved for 18-24 h before feeding on donor calves. For study A, *Ae. aegypti* and *S. calcitrans* were briefly anaesthetised on the day prior to feeding using CO_2_ (*Ae. aegypti*) or chilling (*S. calcitrans*) and 10 insects transferred into a clear-walled pot with a gauze covering. For studies B-D 12 insects were briefly anaesthetised on the day prior to feeding using CO_2_ (*Ae. aegypti*) or chilling (*S. calcitrans*) and placed into cardboard pots with a gauze covering. In studies A-C insects were fed on skin lesions of donor calves. The hair at each feeding site was removed by clipping and the insects, housed in a plastic or cardboard container covered by mesh, held in close contact with the lesion to allow the insects to probe and feed through the mesh. Insects were held in contact with the lesion for approximately 30 sec for interrupted feeding (study A), and for 10-20 min for complete feeding (study B and C).

In studies D and E an artificial membrane feeding system (AMFS) consisting of a Hemotek™ heated reservoir and Parafilm™ membrane was used instead of donor calves. *Aedes aegypti* mosquitoes (study D) or *C. nubeculosus* midges (study E) were fed on defibrinated horse blood (TCS Biosciences Ltd.) spiked with LSDV (10^6^ pfu/ml) that had been purified through a sucrose cushion. In study D, *Ae. aegypti* were fed for 90 sec on the AMFS then fed immediately (within 90 min) on the recipient calves. In study E *C. nubeculosus* were fed for 30 min on the AMFS then incubated for 4 d before being fed on a recipient calf.

Insects were either allowed to blood-feed on recipient calves immediately after feeding on donor cattle or AMFS (studies A and D) or they were incubated for a longer interval (2-4 days) before blood-feeding on recipient cattle (studies B, C and E). After donor or AMFS feeding in studies B, C and E, insects were anaesthetised using CO_2_ and transferred into new cardboard pots. The insects were maintained on 10% sucrose solution (diluted in water) (*Ae. aegypti* and *C. nubeculosus*) or defibrinated horse blood (*S. calcitrans*). Between initial and recipient feeding insects were kept in a temperature-controlled room at ∼25°C, with a 10:14 light/dark cycle. Insects were housed in cardboard/waxed pots with a mesh cover inside a second larger plastic box also covered by mesh. These were kept under a plastic shelter to minimize temperature and humidity fluctuations.

Recipient cattle were prepared for insect feeding by clipping the dorsum of the calves, either side of the midline. The insects, housed in a plastic or cardboard container covered by mesh, were held in close contact with the skin for 10 -20 min. In each of the five studies, recipient feeding occurred over five sequential days, with two pots fed on each calf each day. The feeding sites moved from cranial to caudal each day midline to avoid any area of skin being fed on more than once. After recipient feeding insects were transferred to the laboratory, frozen and kept at -80 °C until processing.

### Collection of additional samples

In addition to insect feeding serum, whole blood, and skin biopsy samples were collected from the calves over the study period. Skin biopsies were collected using a punch biopsy (Kai Pharmaceuticals). The site of biopsy was cleaned with Clinell alcoholic 2% chlorhexidine skin wipes, and the site numbed with 5-8 sec application of a cooling spray. The biopsy was then collected following the manufacturer’s instructions, and placed in a 2mm screw capped tube. Haemostasis of the biopsy site was obtained through manual pressure and application of Celox haemostatic granules.

### Sample analysis

To recover viral and bovine DNA from an insect the whole insect was homogenised in 200µl of homogenisation buffer (calcium free PBS, 1000U Penicillin and Streptomycin (Thermo Fisher) and 4µg/ml Amphotericin B (Thermo Fisher)) using a Tissuelyzer (Qiagen) with metal bead (2 for *S. calcitrans* and *Ae. aegypti* and 1 bead for *C. nubeculosus*) (Dejay Distribution Ltd.) for 30 sec twice at 25Hz, spun at 2,000 rpm for 30 sec and incubated for 30 min at room temperature. DNA extraction was carried out using the MagMax Core extraction kit on the Kingfisher platform (Thermo Scientific) and DNA eluted in 50µl.

Viral DNA was extracted from EDTA blood samples using the KingFisher Flex extraction instrument (Thermo Fisher Scientific) and MagMax Core Extraction kit (A32700; Thermo Fisher Scientific) using Workflow A for DNA extraction from whole blood. The MagMAX_CORE_No_Heat protocol was used as per the manufacturer’s guidelines, with minimal modifications. DNA was eluted in 50 μL of elution buffer.

To determine viral load in the skin, the biopsies were digested by adding 90 μL of proteinase K buffer (4489111, Thermo Fisher Scientific) and 10 μL proteinase K (25530049, Thermo FisherScientific) to the skin biopsy. The sample was incubated for 2 h at 55°C. Tubes were lightly vortexed during incubation and centrifuged briefly. 20 μL of MagMAX Core magnetic beads (Thermo Fisher Scientific) were added to the lysate and mixed. 100 μL of the bead and lysate mixture was transferred to the KingFisher deep well plates (Thermo Fisher Scientific) containing 350 μL of lysis solution and 350 μL of binding solution. Extraction was done as outlined in the MagMAX core extraction kit and the MagMAX_CORE_No_Heat protocol was used to extract DNA from the samples. DNA was eluted in 50 μL of elution buffer.

In study A, analysis of the bovine microsatellite markers present in the bovine blood within the insects was used to identify whether the insect had fed on the donor, recipient or both (methodology described above).

In studies B and C the number of insects that fed on the donor were determined by detection of LSDV genome using PCR. The number of insects that had fed on a recipient was determined by detection of bovine cytochrome B gene. The blood meal from donor would have been digested hence detection of bovine blood meal as an indication of feeding on the recipient was sufficient (Pitzer et al., 2014). The two PCRs were combined in a multiplex PCR. For detection of LSDV genome: forward primer GGCGATGTCCATTCCCTG and reverse primer AGCATTTCATTTCCGTGAGGA was used with probe -ABY CAATGGGTAAAAGATTTCTA QSY. For bovine cytochrome B detection: forward primer – GTAGACAAAGCAACCCTTAC and reverse primer GGAGGAATAGTAGGTGGAC and probe - FAM TTATCATCATAGCAATTGCC MGBNQF was used. Probes were supplied by Thermo Scientific. DNA standards were made by linearising the plasmid LSDV 068 (19AEL6VP; Thermo Fisher) and bovine cytochrome B plasmid (19AEL7QP; Thermo Fisher Scientific) were used to quantify LSDV and bovine cytochrome B respectively. The qPCR conditions were as follows: TaqMan multiplex master mix (4461884; Thermo Fisher Scientific); nuclease free water (10977035; Invitrogen) and 5µl of eluted DNA in a total volume of 20µl was used in the PCR reaction. The cycling conditions included an initial denaturation of 95°C for 20 sec followed by 50 cycles of denaturation at 95°C for 15 sec, annealing and signal capture at 58°C for 1 min.

Study D and E examined transmission of LSDV from an AMFS to a recipient calf. The multiplex PCR described above was used to identify mosquitoes and midges that had acquired LSDV from an AMFS (LSDV detection) and had fed on a recipient calf (bovine cytochrome B detection). The AMFS contained LSDV in horse blood, which would not be detected by the bovine cytochrome B PCR, hence detection of a bovine blood meal as an indication of feeding on the recipient was sufficient.

Detection of LSDV in the blood and skin of the calves was carried out with a quantitative PCR method based on amplification of ORF068 encoding a poly(A) polymerase (small subunit) gene (Tulman et al., 2001, Balinsky et al., 2008). The LSDV Fluorescent Virus Neutralisation Test (FVNT) and partial neutralisation test were carried out as described (Fay et al., 2022) previously. Partial neutralisation was determined by counting the number of fluorescent foci at each dilution using a cut-off of 50 foci per well and converting this into a neutralisation percentage.

### Modelling

The probability that recipient calf *r* became infected (with onset of viremia on day *T_r_* after the initial feed) is given by

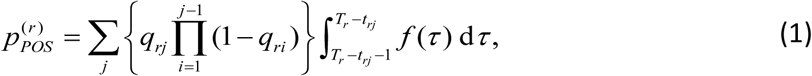

Where

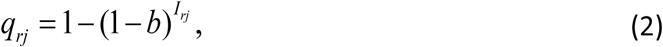

is the probability that recipient *r* became infected on the *j*th feed, *b* is the probability of transmission from insect to bovine, *I_rj_* is the number of infected insects feeding on recipient *r* at feed *j* (Table 1) and *t_rj_* is the time of feed *j* on recipient *r* and *f* is the probability density function for the gamma-distributed latent period (with mean *μ_E_* and shape parameter *k_E_*). The first term in the summation is the probability that the recipient became infected on the *j*th feed (and was not infected in any of the preceding feeds). The second term (the integral) is the probability that, if the animal became infected on the *j*th feed, the onset of viremia would occur on day *T* after the initial feed.

The probability that recipient *r* remained uninfected is given by the probability that transmission did not occur at any of the feeds, that is

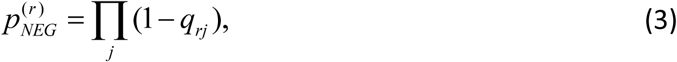

where *q_rj_* is given by equation (2).

### Bayesian methods

Parameters were estimated in a Bayesian framework. The likelihood for the data is

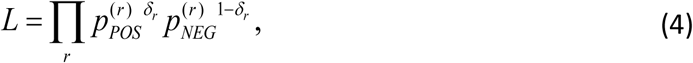

where *δ_r_* is an indicator variable for whether (*δ_r_*=1) or not (*δ_r_*=0) recipient *r* became infected and *p_POS_* and *p_NEG_* are given by equations (1) and (3), respectively. A uniform prior with range (0,1) was used for the probability of transmission from insect to bovine (*b*). Priors for the time to onset of viraemia parameters (*k_E_* and *μ_E_*) were constructed based on analysis of previous challenge experiments(Sanz-Bernardo et al., 2021). Specifically, a gamma prior with mean 3.94 and shape parameter 9.87 was used for *k_E_* and a gamma prior with mean 5.50 and shape parameter 59.9 was used for *μ_E_*.

For *Ae. aegypti* and *S. calcitrans* samples from the posterior distribution were generated using an adaptive Metropolis scheme (Haario et al., 2001), adapted so that the scaling factor was tuned during burn-in to ensure an acceptance rate of between 20% and 40% for more efficient sampling of the target distribution (Andrieu and Thoms, 2008). Two chains of 600,000 iterations were run, with the first 100,000 iterations discarded to allow for burn-in of the chain. The chains were thinned (by selecting every 50th sample) to reduce autocorrelation amongst the samples. Convergence of the scheme was assessed visually and using the Gelman-Rubin statistic provided in the coda package(Plummer et al., 2006, R Development Core Team, 2016) (Plummer et al. 2008) in R (version 4.2.0) (R Core Team 2022).

Because none of the challenges with *C. nubeculosus* resulted in transmission, the posterior distribution for *b* can be computed explicitly. Assuming a uniform (i.e. Beta(1,1)) prior distribution the corresponding posterior distribution is

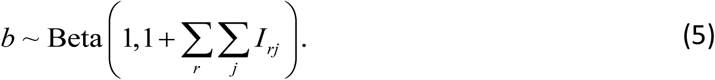

The model for each species was checked using posterior predictive *P*-values. Specifically, the posterior distribution for the model was sampled and the probability that each recipient became infected was computed. Whether or not the recipient became infected after being fed on by infected insects was then simulated, and the observed outcomes were compared to the simulated ones. This procedure was repeated multiple times, and the proportion of samples for which the observed and simulated outcomes matched was computed (i.e., the posterior predictive *P*-value).

### Basic reproduction number

The basic reproduction number (*R*_0_) for transmission of LSDV by each species was computed using the posterior distribution for *b* estimated here (with the distributions for the other parameters remaining the same) as described in Sanz-Bernardo et al. (2021).

## Acknowledgements

The authors would like to express their gratitude to the Insectary Team for their support in providing, rearing, and maintaining the insects. Special thanks also go to the Animal Services team for the animal work.

## Ethical statement

Animal experiments were conducted under license P2137C5BC from the UK Home Office at The Pirbright Institute according to the Animals (Scientific Procedures) Act of 1986 and approved by The Pirbright Institute Animal Welfare and Ethical Review Board.

## Author contributions

Conceptualisation (HM, BSB, CJS, JA, CB, LA, KED, SG, PMB), data curation (HM, BSB, SG), formal analysis (HM, BSB, SG, PMB), funding acquisition (BSB, CJS, JA, LA, KED, SG, PMB), investigation (HM, BSB, NW, IRH, PCF, CGC, AC, IB, CJS, KED, PMB), methodology (HM, BSB, NW, IRH, PCF, CGC, AC, SG), project administration (PMB), resources (CB, LA), software (SG), supervision (CJS, KED, PMB), validation (HM, SG), visualisation (HM, SG, PMB), writing – original draft (HM, SG, PMB), writing – reviewing and editing (HM, BSB, NW, IRH, PCF, CGC, AC, IB, CJS, JA, CB, LA, KED, SG, PMB).

